# Differential laboratory passaging of SARS-CoV-2 viral stocks impacts the in vitro assessment of neutralizing antibodies

**DOI:** 10.1101/2023.07.14.549044

**Authors:** Aram Avila-Herrera, Jeffrey A. Kimbrel, Jose Manuel Marti, James Thissen, Edwin A. Saada, Tracy Weisenberger, Kathryn T. Arrildt, Brent Segelke, Jonathan E. Allen, Adam Zemla, Monica K. Borucki

**Affiliations:** Lawrence Livermore National Laboratory, Computing Directorate, Global Security Computing Division, Livermore, CA, United States of America; Lawrence Livermore National Laboratory, Physical and Life Sciences Directorate, Biotechnology and Biosciences Division, Livermore, CA, United States of America

**Keywords:** SARS-CoV-2, laboratory passage, neutralization, monoclonal antibody, mutation, variant, epitope, binding affinity

## Abstract

Viral populations in natural infections can have a high degree of sequence diversity, which can directly impact immune escape. However, antibody potency is often tested in vitro with a relatively clonal viral populations, such as laboratory virus or pseudotyped virus stocks, which may not accurately represent the genetic diversity of circulating viral genotypes. This can affect the validity of viral phenotype assays, such as antibody neutralization assays. To address this issue, we tested whether recombinant virus carrying SARS-CoV-2 spike (VSV-SARS-CoV-2-S) stocks could be made more genetically diverse by passage, and if a stock passaged under selective pressure was more capable of escaping monoclonal antibody (mAb) neutralization than unpassaged stock or than viral stock passaged without selective pressures. We passaged VSV-SARS-CoV-2-S four times concurrently in three cell lines and then six times with or without polyclonal antiserum selection pressure. All three of the monoclonal antibodies tested neutralized the viral population present in the unpassaged stock. The viral inoculum derived from serial passage without antiserum selection pressure was neutralized by two of the three mAbs. However, the viral inoculum derived from serial passage under antiserum selection pressure escaped neutralization by all three mAbs. Deep sequencing revealed the rapid acquisition of multiple mutations associated with antibody escape in the VSV-SARS-CoV-2-S that had been passaged in the presence of antiserum, including key mutations present in currently circulating Omicron subvariants. These data indicate that viral stock that was generated under polyclonal antiserum selection pressure better reflects the natural environment of the circulating virus and may yield more biologically relevant outcomes in phenotypic assays.

## 1. Introduction

Severe acute respiratory syndrome coronavirus 2 (SARS-CoV-2) has been notably efficient at rapidly evolving resistance to monoclonal antibodies (mAbs) by producing versions of the spike protein with a diversity of amino acid changes that interfere with antibody binding. Importantly, the increased genetic diversity of viral populations that are generated within immunocompromised hosts threatens the efficacy of antibody treatments that are a vital resource for this vulnerable population (1–3). Neutralization assays performed to identify antibody escape mutants generally use a laboratory stock of SARS-CoV-2 strain or a pseudotyped or chimeric virus with a spike protein that represents the wild type Wuhan-Hu-1 strain (3). Passage of viruses in established laboratory cell lines is a convenient way to maintain viral stocks, however the impact of laboratory passage on the genetic diversity of rapidly mutating virus may skew the results of experiments (4–7). While this enables comparison of results between studies, this laboratory stock may not reflect the genetic diversity of circulating strains of the virus.

In this study we passaged a replication-competent vesicular stomatitis virus (VSV) expressing a modified form of the SARS-CoV-2 spike protein (VSV-SARS-CoV-2-S) (8) in three different cell lines and then with or without antiserum selection pressure (Figure 1). Because VSV is an RNA virus and thus has a highly error-prone RNA-dependent RNA polymerase, passage of the spike protein in the VSV backbone rapidly generates mutations in the spike protein during replication (9). We hypothesized that this type of passage would increase the genetic diversity of the viral spike protein and enable the virus to more readily escape antibody neutralization as compared to virus passaged without selection pressure.

**Figure 1.**
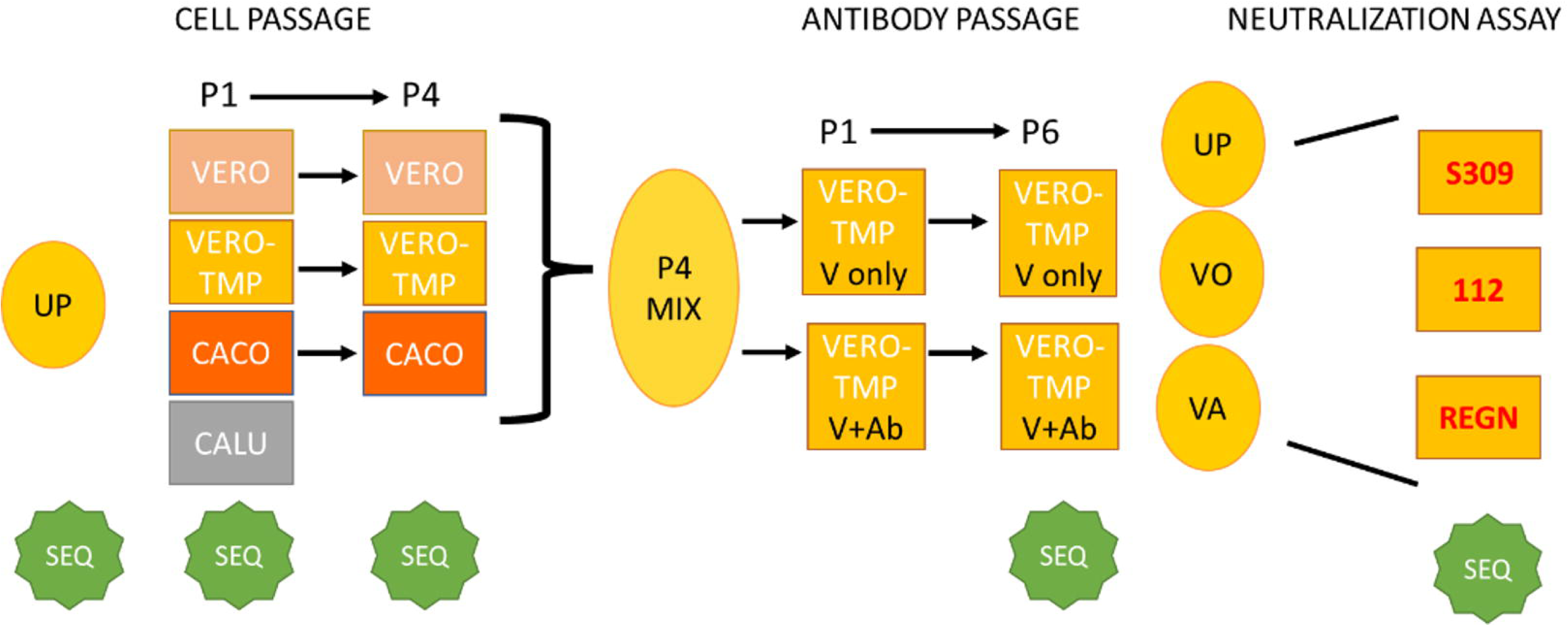
Virus passage and assay workflow. Cell lines and assay components are in boxes, viral inocula are in ovals. UP: unpassaged virus; SEQ: samples were deep sequenced; VERO-TMP: VERO/TMPRSS2; P4 mix: mixture of virus from passage 4 all cell lines; VO: virus only (no antibody selection), VA: virus passaged in the presence of antibody (antiserum from vaccinated individuals).

The SARS-CoV-2 spike protein consists of two subunits, S1 and S2. The S1 subunit binds to a host cell receptor and includes two regions important for antibody evasion, the N-terminal domain (NTD), and the receptor-binding domain (RBD). The S2 subunit is separated from the S1 subunit by a furin cleavage site and functions in virus-cell membrane fusion. To identify mutations associated with passage and antibody escape, the VSV-SARS-CoV-2-S spike gene was deep sequenced prior to passage, after passage in different cell lines, after passage under antiserum selection, and after mAb escape (Figure 1). The mutations that were detected after passage and selection pressure were analyzed using epitope analysis and protein structural modeling.

The cell lines chosen for passage are cell lines commonly used for propagation of coronaviruses that have previously been shown to impact virus mutation (7, 10, 11). Vero and Vero/TMPRSS2 cell lines are highly susceptible to SARS-CoV-2 and generate high viral titers, whereas Calu-3 and Caco-2 cells represent cell types found in biologically relevant tissues, the human respiratory tract and human gastrointestinal tract, respectively, but SARS-CoV-2 may grow to lower titers in these cell lines (7, 12).

## 2. Materials and Methods

### 2.1 Virus

A replicating VSV-SARS-CoV-2-S virus (recombinant virus VSV-SARS-CoV-2-S SΔ21, a gift from Sean Whelan laboratory, Univ. of Washington), which expresses eGFP and has a stop mutation at spike residue 1253 was used for analysis of mutations associated with cell passage and antibody escape (8). Stock VSV-SARS-CoV-2-S virus was generated by passaging once in Vero/TMPRSS2 cells (a gift from Sean Whelan lab, Univ of Washington) at an MOI of 0.05 (as determined by TCID50 assay). The stability of the virus genotype after passage was confirmed via Sanger sequencing and by Illumina sequencing. No changes in the consensus sequence were detected.

### 2.2 Cell lines

The following cell lines were used for passage of the virus: Vero-TMPRSS2 cells, Caco-2 Human intestinal cells (ATCC HTB-37), Vero E6 African Green Monkey kidney cells (ATCC CRL-1586), and Calu-3 Human lung epithelial (ATCC HTB-55). The cell lines were propagated in DMEM media supplemented with 1% Penicillin/Streptomycin and 5 – 10% fetal bovine serum.

### 2.3 Antiserum and monoclonal antibodies

Polyclonal antiserum, Pooled Human Serum Sample, Pfizer Vaccine (NRH-17727) (13), was obtained from BEI Resources. Human Anti-SARS-CoV-2 Spike-RBD Clone S309 mAb and Human Anti-SARS-CoV-2 Spike-RBD REGN10987 mAb (Imdevimab) were purchased from Cell Sciences (Newburyport MA, USA). 2130-1-0114-112 (LLNL-112) is an antibody developed at LLNL (14) derived from COV2-2130 or ciglavimab (15).

### 2.4 Passage on cell lines

Virus was serially passaged four times on three different cell lines to determine whether different mutations are generated in different cell lines. “Unpassaged” stock virus (UP) was used to infect Vero, Vero/TMPRSS2, Caco-2 and Calu-3 cell lines at an MOI of 0.25. After 48h the supernatant was collected and clarified (1000 x g for 5 min) and 100 µl of the supernatant was used for the subsequent passage experiments.

### 2.5 Passage in presence of polyclonal antiserum

To increase the genetic diversity of viral preparations, diluted polyclonal antiserum (Pooled Human Serum Sample, Pfizer Vaccine NRH-17727) was incubated with viral preparations for 1h at 37 °C (Table 1). One hundred microliters of virus-antibody mixtures were used to infect Vero/TMPRSS2 cells in 6-well plates (9). After 48h the two wells with the lowest dilution of antiserum were combined for the next passage. Virus was also passaged without antiserum present to differentiate mutations due to serial passage from mutations due to antibody selection pressure. In each case, virus was serially passaged for a total of six passages.

**Table 1.** Antiserum dilutions used and collected per passage for selection pressure. Three different dilutions of antiserum were used per passage (1-6) in Vero/TMPRSS2 cells. The supernatant from the two lowest dilutions showing CPE and expressing GFP were collected and combined for the subsequent passage. Passage 6 dilutions 1:10 and 1:5 were combined for use in the neutralization assay. GFP: Green fluorescent protein; CPE: Cytopathic effect

| Passage | Dilution 1 | Dilution 2 | Dilution 3 | Dilutions with GFP/CPE | Dilutions collected |
| --- | --- | --- | --- | --- | --- |
| 1 | 1:100 | 1:200 | 1:400 | All | 1 and 2 |
| 2 | 1:50 | 1:100 | 1:200 | All | 1 and 2 |
| 3 | 1:10 | 1:20 | 1:50 | All | 1 and 2 |
| 4 | 1:5 | 1:10 | 1:20 | All | 1 and 2 |
| 5 | 1:5 | 1:10 | 1:20 | All; 1:5-GFP but little CPE | 1 and 2 |
| 6 | 1:5 | 1:10 | 1:20 | All; 1:5-GFP but not CPE | 1 and 2 |

### 2.6 Neutralization assay

To determine whether passage history (cell line and antibody selection pressure) enabled mutant escape of monoclonal antibodies, dilutions of monoclonal antibodies (10 μg/ml or 5 μg/ml) were incubated with 5 × 104 infectious units of virus for 1 hr at 37 °C then applied to monolayers (90% confluence) of Vero/TMPRSS2 cells in 96-well plates in duplicate wells. Supernatant from duplicate wells were pooled for sequence analysis. Three different monoclonal antibodies were tested: S309, REGN10987, and mAb LLNL-112.

### 2.7 RT PCR

RNA was extracted from 140 µl of culture supernatant using the Qiagen QIAamp Viral RNA Mini Kit. Viral RNA was amplified and sequenced using a version of the ARTIC protocol, nCoV-2019 sequencing protocol (https://www.protocols.io/view/ncov-2019-sequencing-protocol-v3-locost-bp2l6n26rgqe/v3, accessed 12/22/2022). Because the spike gene may mutate rapidly, the spike gene was amplified using two ARTIC primer sets, v3 and 4.1 to increase the chance of mutations being detected. All of the samples with exception of cell passage 1 and 4 (Figure 1) were amplified in duplicate PCR reactions, thus for each of these samples 8 different PCR reactions were performed (each ARTIC primer set requires 2 different PCR reactions and two ARTIC primer versions were used in replicates). The RT reaction was performed using the Luna (NEB). The cDNA was amplified in a 25 µl PCR reactions using 6 µl cDNA and 2X Master mix (NEB). The PCR cycling program was 98 °C for 30 seconds, then 35 cycles of 98 °C for 15 seconds, 65 °C for 5 minutes. A negative PCR control for each PCR set was prepared using PCR grade water as a template. Negative control PCR products were combined from different PCR batches and was sequenced as a negative control. The PCR products from each reaction were pooled then were purified using AMPureXP beads (Beckman Coulter, Brea, CA, USA). A negative control (no template control, NTC) was prepared by pooling the NTC PCR products from multiple runs from each primer set.

### 2.8 Sequencing

A total of 40 µL of pooled v3 and v4.1 ARTIC amplicons were purified by adding 0.8X of Ampure XP magnetic beads (Beckman Coulter) to each pooled amplicon sample. The PCR products were purified following the standard manufacturer’s instructions and resuspended in 22 µL of water. The purified PCR products were quantified using the Qubit fluorimeter (ThermoFisher) and 100 ng of each sample was input into the Illumina DNA Prep library kit (Illumina). The small amplicon manufacturer’s instructions were followed for the library kit and completed libraries were quantified and verified on the Tapestation 4200 instrument (Agilent). The completed libraries were diluted to 4 nM, pooled, and further diluted to a loading concentration of 8 pM. The libraries were sequenced on the Illumina MiSeq instrument using a v2 2x250 sequencing kit. Raw sequencing data has been uploaded to the NCBI under BioProject PRJNA978027.

### 2.9 Bioinformatic analysis

Variants were called using Mappgene, (https://github.com/LLNL/mappgene) a SARS-CoV-2 variant calling pipeline for high performance computing environments (HPC) (16). Mappgene employs BWA-MEM (17) for read alignment, iVar 1.3.1 for read trimming/filtering/variant calling, and LoFreq v2.1.5 to call variants (18).

### 2.10 Epitope analysis

The epitope regions for three SARS-CoV-2 neutralizing antibodies and the hACE2 receptor binding motif (RBM) were modeled to determine if epitopes are in close proximity or overlapping the RBM, and if the locations of the observed mutations might impact binding (Figure 2). The coordinates for the S309-RBD complex comes from PDB entry 7R6W; the Regeneron REGN10987-RBD complex from PDB entry 6XDG (chains A and C make up the REGN10987 light and heavy chain, respectively, Fab complexed with RBD [chain E]); and the LLNL-112-RBD complex comes from a model structured generated at Lawrence Livermore National Laboratory (LLNL) which was in turn produced from the COV2-2130 complex with RBD coordinates in PDB entry 7L7E. The RBM was determined from the human ACE2-SARS-CoV-2 RBD complex coordinates in PDB entry 6M0J. The amino acids making up the epitopes and RBM were determined from the listed coordinate sets as described and the region made up of these residues is outlined on the RDB molecular surface from PDB entry 7R6W; the epitopes and RBM were outlined on a common RBD molecular surface for ease of comparison.

**Figure 2.**
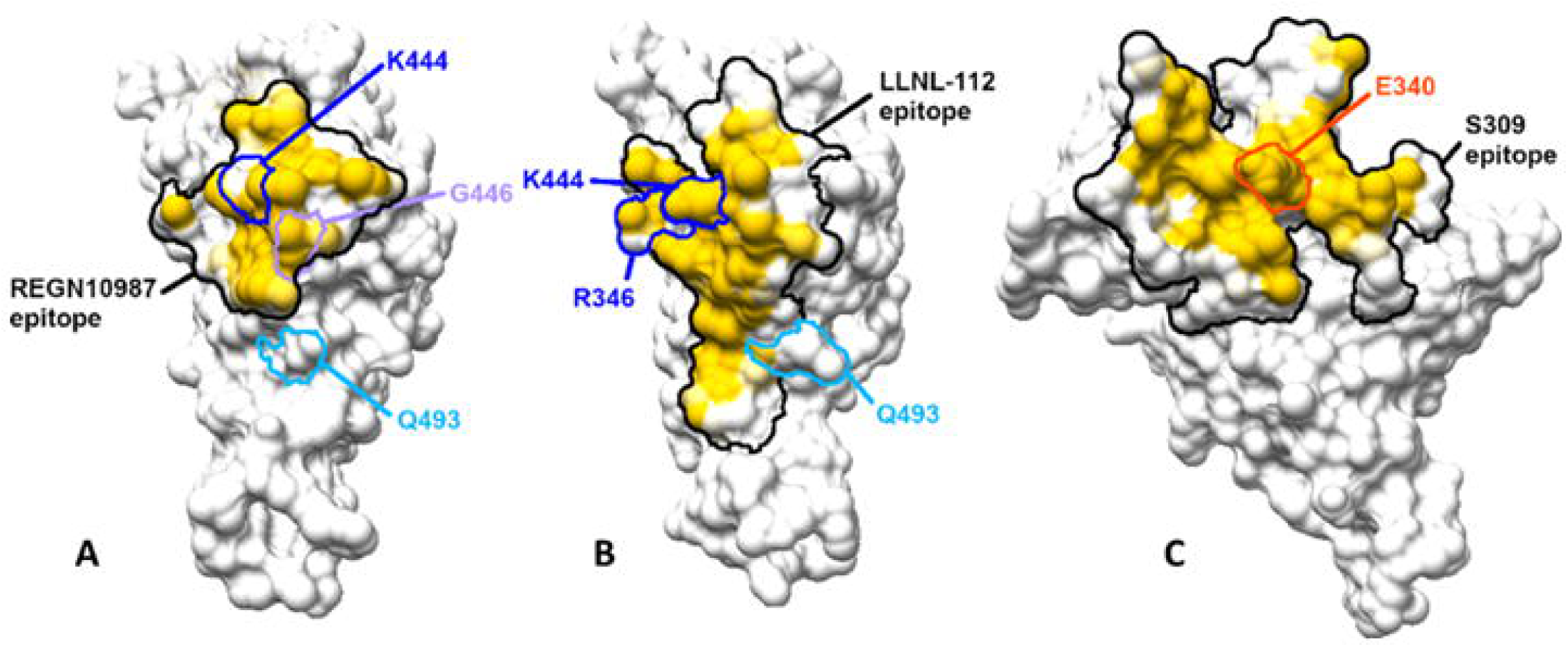
Molecular surfaces for the epitope regions for neutralizing antibodies and amino acids that are prevalent in genotypes that escaped antibody neutralization. The epitopes are outlined and colored by degree of buried surface in the antibody-antigen complex. The black outlined region encloses all atoms in any residue that has any atom within 5 Å of any atom of the antibody and contributes to the molecular surface. A) REGN10987 epitope mapping shows two amino acid locations at which escape mutations occur, 444 and 446, are in the epitope; K444 and G446 have a high buried surface area in the complex with REGN10987, thus mutations at these positions could have high impact on the intermolecular interaction. B) LLNL-112 epitope mapping shows the location of escape mutations (346 and 444) occur in the epitope, and K444 is highly buried in the complex. C) Molecular surfaces for the epitope regions for S309 and amino acids prevalent in genotypes that escape S309, E340A and R346S. E340A is in the center of the epitope and has significant buried surface in the complex. These figures were generated using Chimera (21) and GIMP (The GIMP Development Team, 2019. GIMP, Available at: https://www.gimp.org).

### 2.11 Protein structural analysis

To assess if the escape mutations increased ability to evade neutralizing antibodies, we applied our Structural Fluctuation Estimation (SFE) method (19) to predict their impact on different mAbs in complex with the Wuhan as well as the Omicron strains. To make more complete comparison, the estimations of binding affinity changes (ΔΔG - predicted changes in binding energy upon introduced mutations) were also calculated for the RBD-ACE2 complexes (19). SFE starts with the construction of structural models for each of the RBD-mAb and RBD-ACE2 complexes. For each constructed complex we performed minimization and relaxation using disparate approaches: Rosetta (20), Chimera (21), and GROMACS (22) steepest decent and conjugant gradient minimizations, and molecular dynamics trajectories from GROMACS simulations. This procedure as applied to each of constructed complexes allowed us to sample an extended number of structural conformations, use them to perform “forward” and “reverse” ΔΔG calculations using Flex ddG (23), remove outliers, average results, and report with higher confidence final energy change estimates. By computing ΔΔG’s across many conformations of each constructed complex, we reduced uncertainties in the structural modeling and estimated natural conformational fluctuations within the complexes.

## 3. Results

### 3.1 Virus passage on different cell lines

To determine the effect of host cell line on viral diversity, virus was passaged four times on different cell lines: Vero, Vero/TMPRSS2, Caco-2 and Calu-3. Unpassaged stock virus (UP) was used to infect each cell lines at an MOI of 0.25. After 48 h the supernatant was collected and 100 μl of the supernatant was used to for the subsequent passage experiments. This was repeated for a total of 4 passages per cell line with the exception of Calu-3 cells. Calu-3 cells had no cytopathic effect (CPE) evident after 48 h post infection and few cells expressing GFP per field after the first passage. No GFP expression or CPE was observed in passage 2 of the Calu-3 cells. The 2nd passage was repeated a total of 3 times with the same result. Calu-3 cells over expressing human ACE-2 receptor was obtained from BEI Resources (Clone 2B4, NR-55340) and passage was tried using this cell line with no evidence of CPE or GFP expression, although inoculation of Calu-3 passage 1 (P1) supernatant onto Vero/TMPRRS2 cells did yield strong GFP expression.

After 4 passages in Vero, Vero/TMPRSS2, or Caco-2 cells, supernatant from the final passage in each cell line was titered by TCID50 and equal amounts of passaged virus (2 x 105infectious units) were combined into a single inoculum for subsequent passages under antibody selection pressure.

### 3.2 Virus passage with antibody selection pressure

To understand how antibody selection pressure can modify the mutational profiles, this “Cell-passaged” inoculum was used to infect Vero/TMPRSS2 cells at an MOI of 0.1 in the presence of polyclonal antiserum (“Virus and antibody”, VA) or in media without the presence of polyclonal antiserum (“Virus only”, VO). For passage in the presence of sera, three different dilutions of antisera were used per passage, and supernatant from the two lowest dilutions showing CPE and expressing GFP were collected and combined for the subsequent passage. Early passages showed that highly diluted sera did not inhibit viral growth thus sera concentrations were increased in subsequent passages (Table 1).

### 3.3 Genetic analysis of mutations associated with cell passage and antiserum selection pressure

To identify mutations associated with passage in cell lines and in the presence of antiserum, the spike gene within the VSV backbone was amplified using multiplexed PCR (ARTIC V3 protocol) and sequenced using the Illumina platform. The UP stock and passaged samples (VO, VA) were PCR amplified and sequenced in duplicate to increase confidence in detection of low frequency mutations. The ARTIC primer sets are designed to span the genome (versus just the spike gene), and the highly multiplexed primer sets amplified 97% of the spike gene present in the VSV vector and did not appear to be negatively impacted by the presence of primers designed to amplify other regions of the SARS-CoV-2 genome. Although the RT-PCR amplicons often produced many bands larger than the target size of 400 bp, sequencing coverage was excellent with >95% of the spike gene having coverage of at least 1000 x depth (Supp data Fig 1). None of the negative control samples showed bands of any size on the PCR gels. The first sequencing run had a maximum of 11 reads mapped to any nt position for the negative PCR control (NTC), the second run had no mapped reads from the NTC.

Analysis of the deep sequence data revealed no mutations were detected at >1% frequency in the UP stock (sequenced in duplicate). A high frequency mutation, R685S, was detected in Vero P1 at 5.2% and at 83.8% frequency in Vero passage 4 (P4) (Table 2). Interestingly, VSV-GFP-SARS-CoV-2-S Δ21 with the R685S mutation has previously been isolated from WT stock (24). Four mutations were detected in Vero/TMPRSS2 P4 and these mutations were enriched in later samples, often at much higher frequencies: L48S, T76I, H655Y, and I1081V (Tables 2 and 3, Supp Table 3). Caco-2 P4 had only one mutation detected at >1%, I468T detected at 2.2% frequency (Table 2).

**Table 2.**
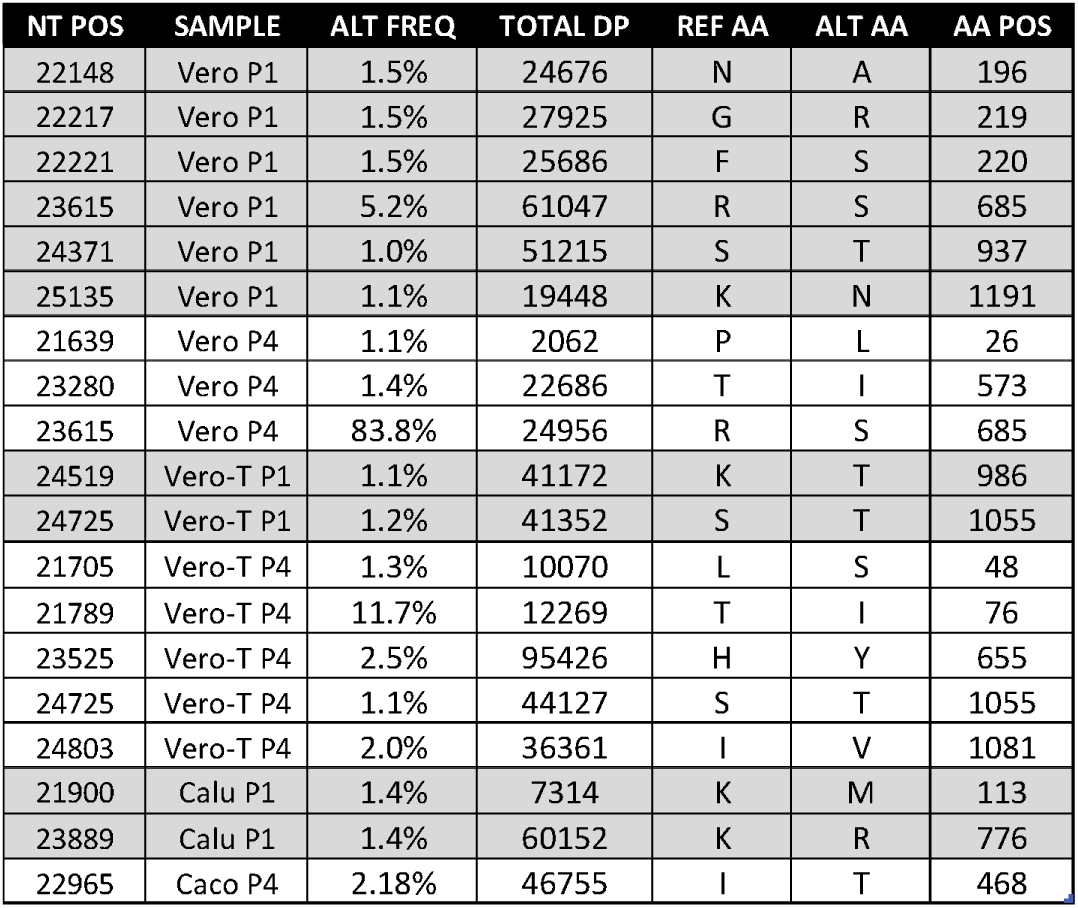
Mutations detected after passage in cell lines. Abbreviations Vero-T: Vero/TMPRSS2; NT POS: Nucleotide position in the reference sequence Wuhan-Hu-1 (NC_045512.2); AVG FREQ: Average frequency; TOTAL DP: Depth of sequence read coverage for the nt pos; REF AA: Amino acid in the reference sequence; ALT AA: Amino acid in the sample sequence; AA POS: Residue number in the spike gene.

| NT POS | SAMPLE | ALT FREQ | TOTAL DP | REF AA | ALT AA | AA POS |
| --- | --- | --- | --- | --- | --- | --- |
| 22148 | Vero P1 | 1.5% | 24676 | N | A | 196 |
| 22217 | Vero P1 | 1.5% | 27925 | G | R | 219 |
| 22221 | Vero P1 | 1.5% | 25686 | F | S | 220 |
| 23615 | Vero P1 | 5.2% | 61047 | R | S | 685 |
| 24371 | Vero P1 | 1.0% | 51215 | S | T | 937 |
| 25135 | Vero P1 | 1.1% | 19448 | K | N | 1191 |
| 21639 | Vero P4 | 1.1% | 2062 | P | L | 26 |
| 23280 | Vero P4 | 1.4% | 22686 | T | I | 573 |
| 23615 | Vero P4 | 83.8% | 24956 | R | S | 685 |
| 24519 | Vero-T P1 | 1.1% | 41172 | K | T | 986 |
| 24725 | Vero-T P1 | 1.2% | 41352 | S | T | 1055 |
| 21705 | Vero-T P4 | 1.3% | 10070 | L | S | 48 |
| 21789 | Vero-T P4 | 11.7% | 12269 | T | I | 76 |
| 23525 | Vero-T P4 | 2.5% | 95426 | H | Y | 655 |
| 24725 | Vero-T P4 | 1.1% | 44127 | S | T | 1055 |
| 24803 | Vero-T P4 | 2.0% | 36361 | I | V | 1081 |
| 21900 | Calu P1 | 1.4% | 7314 | K | M | 113 |
| 23889 | Calu P1 | 1.4% | 60152 | K | R | 776 |
| 22965 | Caco P4 | 2.18% | 46755 | I | T | 468 |

**Table 3.**
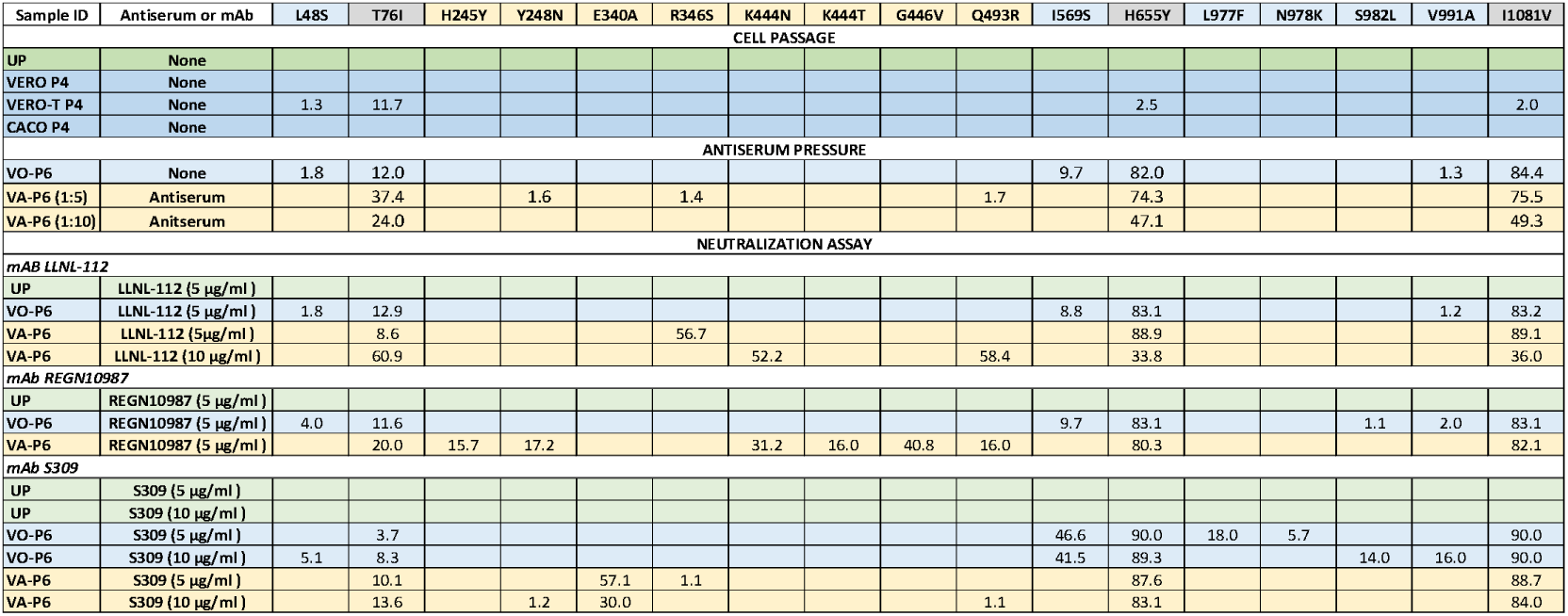
Comparison of variant frequencies associated with cell culture passage, passage under antiserum selection, and mAb selection. Mutations that are present in at least 1 sample at > 5% are shown and that are present in antiserum passaged samples and/or neutralization data are shown (see Suppl Table 1 for mutations present at 1% or greater). Note cell passaged samples from Vero and Caco-2 cells did not have mutations that persisted and are not shown. Antiserum passaged samples (VA-P6) have serum dilution (1:5 or 1:10) shown in parentheses. Samples that were tested with the neutralization assay are denoted by the mAb name, the viral inoculum (UP, VO-P6, or VA-P6), and the mAb concentration (5 µg/ml, 10 µg/ml). Data from virus only passaged samples (no antibody selection) are shown in blue cells and data from virus passaged under antiserum or mAb selection are in yellow shaded cells. Top row cells shaded grey signify columns with data from both virus only and antibody selection. With the exception of Vero P4, Vero-T P4 and Caco P4, all samples were processed (PCR and sequencing) in duplicate; the percent frequency is an average of the two values.

After pooling equal amount of virus (P4) from the three cell lines (Fig 1), the virus was further passaged six times in Vero/TMPRSS2 cells either with or without the presence of antiserum. Viral RNA from passage 6 was PCR amplified in duplicate and subjected to deep sequencing. Table 3 shows the location and frequency of the mutations detected.

The six passages with or without antiserum selection pressure induced high frequency mutations in three residues, T76I, H655Y, and I1081V, with H655Y and I1081V appearing to be linked due to these mutations always being detected together and at the same approximate frequency. These mutations were first detected in Vero/TMPRSS2 P4. Interestingly, virus passaged six times without antiserum selection pressure (VO-P6) produced a high frequency mutation, I569S that continued to persist at high frequency exclusively in virus only samples, and was not detected in samples passaged in the presence of antiserum or mAbs (Table 3, Supp Table 3). Alternatively, the three mutations that were located in the RBD were detected only in samples passaged in the presence of antiserum (VA-P6) or mAbs.

### 3.4 Mutations associated with escape from mAb neutralization

To determine whether the mutations that occurred during passage with antiserum selection pressure could confer resistance to mAbs designed to treat COVID infections, supernatant from UP, VA-P6, and VO-P6 were assayed for mAb escape using neutralization assays and deep sequencing (Figure 1). mAbs (5 μg/ml and 10 μg/ml) were incubated with 5 × 104 infectious units of virus for 1 hr at 37°C then applied to Vero/TMPRSS2 cells. Two days post infection cell monolayers were checked for GFP expression and scored as negative (fewer than 10 cells with GFP expression), weakly positive (multiple small patches of GFP), positive (large patches of GFP), or strongly positive (majority of the monolayer showing GFP expression) (Figure 2, Table 4). Inocula containing no mAb (virus only) were used as positive controls and inocula containing antibody but no virus were used as negative controls.

**Table 4.** Neutralization assay comparison of mAb escape by unpassaged and passaged virus. Viral escape was measured by GFP expression 48 h post infection. Note: Wells scored as “+/-“ were considered as negative for mAb escape. Data from deep Illumina sequencing of the samples were used to identify mutations that were associated with mAb escape. These mutations were compared to the mutations that occurred during passage in cell lines and under antiserum selection to understand if the mutations originated during passage or appeared to arise de novo during mAb escape (Table 3). Epitope models were used to visualize the placement of the mutations on the RBD and within the receptor binding motif (RBM) (Figure 2).

| mAb conc.<br>(ug/ml) | Control<br>no IgG | LLNL-112 |  | S309 |  | REGN10987 |  |
| --- | --- | --- | --- | --- | --- | --- | --- |
|  |  | 5 | 10 | 5 | 10 | 5 | 10 |
| No Virus | - - | - - | - - | - - | - - | - - | - - |
| UP | +++ +++ | - - | - - | +/- +/- | +/- +/- | - - | - - |
| VO.P6 | +++ +++ | - - | - - | ++ ++ | ++ ++ | - - | - - |
| VA.P6 | +++ +++ | ++ ++ | - +/- | + +++ | + + | ++ ++ | - - |

Escape from neutralization by mAbs was significantly impacted by passage of the virus. UP virus was neutralized by all three of the mAb’s and VO-P6 was neutralized by 2 of the 3 mAbs (Table 4), whereas virus passed in the presence of antiserum, VA-P6, was able to escape neutralization by all three mAbs at a concentration of 5 µg/ml. Supernatant was collected from wells showing GFP and from wells from the same mAb concentrations that did not have GFP. RNA was extracted from the samples (each sample was collected as a pool of the two replicate wells) and the spike gene was amplified using ARTIC protocol for RT-PCR. Each sample was PCR amplified and sequenced in duplicate to increase confidence in rare variant detection.

The three mutations prevalent in the passaged virus (VO-P6 and VA-P6), T76I, H655Y, and I1081V, were maintained at high frequency regardless of whether the virus escaped neutralization but none of these mutations were detected in the UP samples. Similarly, the I569S mutation was detected exclusively in neutralization samples derived from VO-P6. The frequency of I569S increased approximately 4 fold in the S309 samples, the only mAb for which neutralization escape was detected for VO-P6, but not for REGN10987 or LLNL-112 (Table 3). Multiple high frequency mutations K444N, K444T, G446V, and Q493R were detected in REGN10987 neutralization assay samples derived from VO-P6 (5 µg/ml) (Table 3). All of these mutations have been described previously as affecting neutralization by polyclonal antiserum and mAbs (25, 26). K444T and G446V were detected at the same frequency (16%) and visual inspection of mapped sequencing reads revealed these mutations often co-occur and may be linked.

Two high frequency mutations were detected in the N terminal domain (NTD) of REGN10987 sample VA-P6 (5 µg/ml). These mutations, H245Y (15.7%) and Y248N (17.2%), are located within the NTD antigen supersite (2, 27, 28). REGN10987 is not known to bind the NTD, thus it is possible these mutations are part of genotypes that have additional mutations that impact REGN10987 binding such as K444T and Q493R. Both of these mutations occur in REGN10987 VA-P6 (5 µg/ml) sequence data at approximately the same frequency as H245Y and Y248N. Visual examination of mapped sequencing reads verified that the 245 and 248 mutations are linked, however, most of the Illumina reads were too short to determine if 444 and 493 mutations are linked.

Significant GFP expression was detected in all wells of the mAb S309 neutralization assay except for those of the UP inoculum (Table 4, Supp Fig 2). The mutations associated with escape from S309 differed between VO-P6 and VA-P6 with no high frequency mutations detected in RBD for sequences from VO-P6, but an E340A mutation was detected at 57% and 30% frequency in the sequences from 5 µg/ml and 10 µg/ml wells. Additional mutations at sites associated with mAb, R346S and Q493R were also detected at 1% in sequence data derived from the 5 µg/ml and 10 µg/ml wells. Interestingly sequence data from VO-P6 wells showed three high frequency mutations in the S2 spike subunit, L977F, S982L, and V991A, detected at 18%, 14%, and 16%, respectively. Mutations at two of these residues, S982 and V991, may impact the positioning of the RBD and thus impact infectivity (29–31).

LLNL-112 mAb had a neutralization pattern similar to that of REGN10987 with UP and VO-P6 neutralized by the mAb but escape observed with VA-P6 (Tables 3 and 4). Deep sequencing data showed that three RBD mutations were present in the viral RNA amplified from the 5 µg/ml and 10 µg/ml mAb wells. The R346S mutation was detected in the 5 µg/ml mAb wells at 57% frequency, whereas the 10 µg/ml wells produced different mutations, K444N at 52% frequency, and Q493R at 58% for 10 µg/ml. Notably, the 10 µg/ml wells were scored as negative for viral escape because fewer than 10 GFP positive cells were present, however, the few cells expressing GFP were able to produce enough virus to be detectable by RT-PCR.

### 3.5 De novo mutations in RBD

Many of the mutations that were detected in the RBD at high frequency after mAb selection were not detected in the viral inoculum used in the neutralization assay, even when examining the samples using deep sequencing (detection of mutations present at less than 1% frequency). In particular, mutations E340A, R346S, K444N, K444T, G446V, and Q493R were detected at high frequencies ranging from 16 to 58% in sequences generated from mAb challenged with VA-P6 (Table 3). Deep sequence analysis of VA-P6 revealed that only R346S was present in VA-P6 at 1.4%, whereas the other five mutations were not detected at even very low frequency and thus appear to have been generated de novo, even in the case of VA-P6 inoculum with 10 µg/ml LLNL-112 mAb which had only a few cells expressing GFP. Further examination of the sequence data from the six mutations showed that in each case the amino acid change was due to a single nucleotide mutation, thus requiring a single replication error to generate the amino acid change. These data suggest that RBD escape mutants can rapidly arise de novo and be enriched to become high frequency or consensus mutations.

### 3.6 Protein structure analysis of mutations

The high frequency mutations associated with cell passage, antiserum selection, or antibody escape were modeled to assess potential impact on protein function and antibody binding. Figure 3 shows the relative positions of a subset the mutations that arose during passage. Notably, although H655Y and I1081V appear to be linked, these mutations are not co-located on the spike trimer. Although previous studies have shown that H655Y is frequently detected during passage and in variant genotypes (11, 16, 32), the co-occurrence of I1081V has not been documented. Similarly, mutation I569S was selected for during virus only passage but has not been described in the literature. Figure 3 shows the position of I569S is not adjacent to the RBD on the trimer relative to the RBD. Unfortunately, the residues from the furin cleavage site (681-685) are missing in all experimentally solved spike structures deposited in PDB. Judging from the approximate location of this site based on predicted models, residue 569 is quite far from this site and it is unlikely that I569S may directly affect the cleavage site.

**Figure 3.**
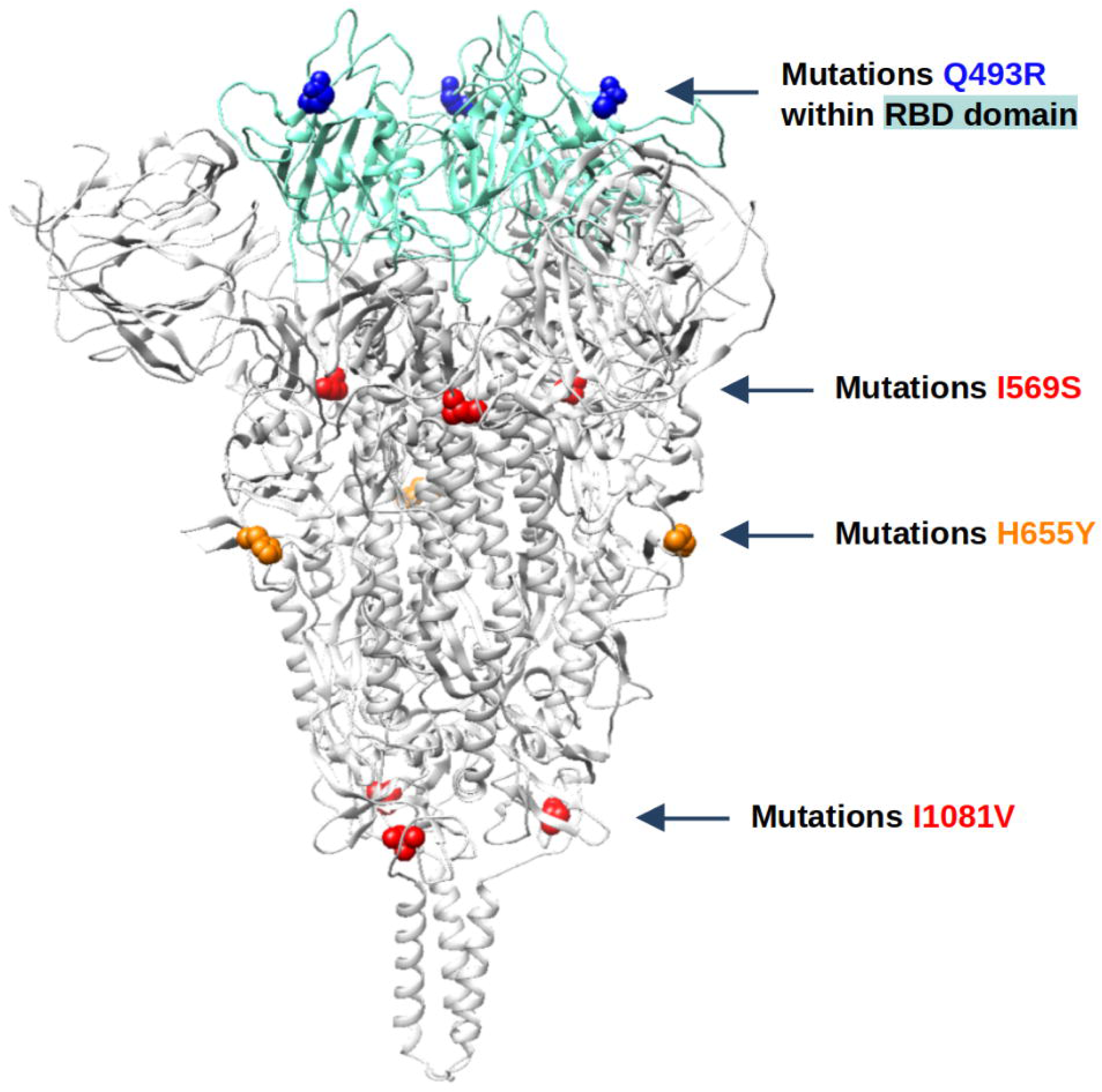
Trimer assembly of the Spike protein with select mutations

Protein structural models of RBD-mAb complexes were constructed to evaluate binding affinity changes of mutations observed in the RBD domain and confirm whether the mutations that occurred during passage with antiserum selection pressure could confer resistance to mAbs (Table 5). A higher positive value for a ΔΔG scores indicates that the RBD mutant is weakening (or breaking) the binding of the mAb. The predicted changes in ΔΔG agree with observed mutant frequencies in 24/27 cases (85%). Indeed, the estimated ΔΔG values show an increase in all positions with mutations when mutations are within RBD-mAb interface (distances between RBD and mAb are below 7 Ångstroms). Thus, the results from the ΔΔG calculations (Table 5) support the hypothesis that the mutations are in fact antibody escape mutations.

**Table 5.** Comparison of VA-P6 mutation frequencies associated with passage under mAb selection with predicted binding affinity changes (ΔΔG) induced by observed mutations. The percent frequency of each mutation (from Table 3) is shown in the yellow-colored rows. In each ΔΔG row (in orange) the distances (Ångstroms) between the closest atoms from the RBD mutation and mAb interface residues are provided in parentheses.

|  | E340A | R346S | K378E | L390F | K444N | K444T | G446V | T478K | Q493R |
| --- | --- | --- | --- | --- | --- | --- | --- | --- | --- |
| **LLNL-112** |  |  |  |  |  |  |  |  |  |
| LLNL-112_AB_5 |  | 56.7% |  |  |  |  |  |  |  |
| LLNL-112_AB_10 |  |  |  |  | 52.2% |  |  |  | 58.4% |
| LLNL-112 $\Delta\Delta G$<br>(RBD-mAb distance) | 0<br>(9.6) | 1.4<br>(2.1) | 0<br>(25.1) | 0<br>(30.7) | 2.3<br>(2.7) | 2.9<br>(2.7) | -0.7<br>(2.9) | 0<br>(16.7) | 0<br>(3.5) |
| **REGN10987** |  |  |  |  |  |  |  |  |  |
| REGN10987_AB_5 |  |  |  |  | 31.2% | 16.0% | 40.8% |  | 16.0% |
| REGN10987 $\Delta\Delta G$<br>(RBD-mAb distance) | 0<br>(13.2) | 0<br>(3.0) | 0<br>(20.5) | 0<br>(32.4) | 0.5<br>(3.3) | 0.7<br>(3.3) | 0.6<br>(2.7) | 0<br>(31.1) | 0<br>(11.9) |
| **S309** |  |  |  |  |  |  |  |  |  |
| S309_AB_5 | 57.1% | 1.1% |  |  |  |  |  |  |  |
| S309_AB_10 | 30.0% |  |  |  |  |  |  |  | 1.1% |
| S309 $\Delta\Delta G$<br>(RBD-mAb distance) | 3.4<br>(2.8) | 2.9<br>(4.8) | 0<br>(19.6) | 0<br>(14.9) | -0.2<br>(5.2) | -0.2<br>(5.2) | 0<br>(9.6) | 0<br>(37.6) | 0<br>(19.1) |

## 4. Discussion

In this study, we determined that VSV/SARS-CoV-2 stock was made more genetically diverse by passage under antiserum selective pressure, and that this passaged viral stock was able to more readily escape monoclonal antibody neutralization as compared to unpassaged stock or viral stock passaged without selective pressures. Deep sequencing detected the presence of multiple mutations associated with antibody escape and enhanced transmissibility in the VSV-SARS-CoV-2-S virus that had been passaged in the presence of antiserum. Analysis of mutations detected in the RBD using epitope and protein structural models indicate that these mutations are a result of antibody escape. These data indicate viral stock that was generated with limited polyclonal antiserum selection pressure may yield more biologically relevant outcomes of phenotypic assays.

In this and other studies (9, 33) the error prone RNA dependent RNA polymerase of the VSV genome enabled rapid generation of escape mutations. Although the mutation rate of SARS-CoV-2 is lower than that of other RNA viruses such as VSV that lack the 311 exonuclease proofreading mechanism found in coronaviruses (34), the escape mutants observed by this and other studies reflect mutations detected in natural infections (9, 33). This can be explained by the large viral populations generated in natural infections and the enormous scope of the CoVID-19 pandemic. A SARS-CoV-2 intrahost population is estimated to consist of over 10^9^ copies of viral RNA (35) and this large viral population enables many different mutations to be generated within a single infected individual thus increasing diversity of mutations generated in natural infections (9). Thus, the rapid mutation rate of the VSV polymerase allowed us to sample a larger number of mutations using a smaller population and a shorter time scale, which made it possible to rapidly identify escape mutations.

To determine the impact of cell type on the genetic diversity of viral populations, we first applied selective pressure by passing the viral stock through three different cell lines Deep sequencing data indicated that passage in these cell lines induced relatively few mutations present at >1%, and only one mutation, I468T detected in Caco-2 cells P4, was detected in the RBD (Table 1).

Many of the mutations that were detected after passage in cell lines had been described in prior studies. For example, a high frequency mutation, R685S, detected Vero cells P1 at 5.2%, and enriched to 83.8% frequency by P4, was previously isolated from VSV/SARS-CoV-2 stock (24). This mutation occurs at the S1/S2 furin cleavage site, and has also arisen previously during passage of SARS-CoV-2 virus in Vero cells (36).

Some expected mutations such as loss of the furin cleavage site during passage in Vero cells and mutation at known immune evasion site E484 were not seen (4, 32). This could be due to the relatively limited number of passages in each cell line, characteristics of the antiserum used for selection pressure, or the size and composition of the viral population used for the passage experiments. It is possible that more extensive passage in more diverse cell lines such as cell lines derived from other organs and other species might yield a more diverse set of mutations (11).

A subset of mutations that arose during cell line passage persisted during six subsequent passages in Vero/TMPRSS2 cells, with or without antiserum selection pressure (Table 3). All of these persistent mutations were found at >1% frequency only in Vero/TMPRSS2 P4, and none of mutations generated by passage in the other two cell lines, Vero or Caco-2, persisted. Four of these mutations, L48S, T76I, H655Y, and I1081V were detected in sequences from virus passage with or without the presence of antibody and escape mutant samples (Table 3). The mutations at residues H655Y and I1081V were detected in the same samples and at similar frequencies, thus appear to be linked. Among the four mutations consistently detected in sequences from virus passage both with or without the presence of antibody, mutations T76I and H655Y are present in variant of concern genotypes. Both these mutations may aid in immune escape and infectivity, however, they are also associated with passage in cell culture (11, 37, 38), thus, the role of these mutations in mAb escape is unclear.

Each of the three mAbs used in the neutralization assays was designed to neutralize SARS-CoV-2 virus but differed in efficacy according to differences in variant genotypes that emerged as the pandemic progressed (3). Commercially available mAbs REGN10987 (Imdevimab) and S309 (Sotrovimab) had been found to be sensitive to different mutations (3), and mutations in Omicron subvariants have rendered both REGN10987 and S309 ineffective or weakly active (39).

Neutralization patterns were similar for REGN10987 and LLNL-112, with both of these mAbs blocking viral replication for UP and VO-P6; only VA-P6 escaped and generated significant infection in the 5 µg/ml wells. The mutations generated during viral replication occurred at RBD residues associated with polyclonal antiserum and mAb escape (25, 26), and within or adjacent to the known epitopes of the antibodies (3) (Fig 2). Mutations at these residues (346, 444, 446, and 493) have been detected in variants of concern including Omicron sublineages (40).

Compared to REGN10987 and LLNL-112, mAb S309 was less effective in blocking the virus present in the VO-P6 and VA-P6 inoculum from entering cells but did block the viral population in the UP inoculum from productively replicating in cells. Interestingly, despite the presence of extensive GFP expression in VO-P6 wells, no high frequency mutations were detected in the RBD, thus breakthrough may have been driven by mutations in other regions of the spike protein that increased viral infectivity. Mutations S982L and V991A may impact infectivity (29, 31) and were detected only in the VO-P6, 10 µg/ml mAb sample, however, the viral sequences derived from the VO-P6, 5 µg/ml mAb sample had different S2 subunit mutations, L977F and N978K. Additionally, the I569S mutation is greatly enriched in the VO-P6 samples as compared to VO-P6 samples from REGB10987 and LLNL-112 (Table 3). It is unclear if this mutation plays a role in infectivity or immune evasion in vitro or in vivo; a query of outbreak.info (41) shows that this mutation is rarely is detected in human sequences. S309 escape by VA-P6 appears to be driven by E340A mutation was detected at high frequency. These data agree with previous studies that identify mutation E340A as significantly impacting neutralization efficacy of S309 (42, 43), and epitope analysis indicates residue 340 is buried within mAb S309 epitope (Fig 2).

For mutations observed in the RBD domain of the spike protein we constructed structural models of RBD-mAb complexes to evaluate binding affinity changes and confirm whether the mutations that occurred during passage with antiserum selection pressure could confer resistance to mAbs. A general grouping of antibody escape mutations can be based on their direct or indirect contribution to the binding event. For example, (1) those that directly interfere with the binding (changes within the interfaces) such as 346 or 444, and (2) those that introduce some changes in local structure conformations outside the Ag-Ab interfaces that impede the antibody’s access. In the former case such mutations may work in conjunction with some additional mutations. Results from protein structural analysis indicate that combinations of mutations enabled antibody evasion: LLNL-112 was escaped by R346S and K444N mutations, REGN10987 escaped by K444N/T and G446V, and S309 binding escaped by E340A and R346S. Mutation Q493R co-occurs with other spike protein mutations and may belong to the second category – indirect impact on the binding. This assumption is supported by the following four observations: (1) Q493R is one of the major mutations observed in the Omicron variant of COVID-19 that increased viral binding to ACE2 and knocked-out many known, circulating antibodies (19), however in all tested RBD-mAb interactions a single-point mutation Q493R alone is not affecting binding: ΔΔG = 0.0, (2) Q493R mutation is outside the RBD interfaces with REGN10987 (11.9 Å) and sotrovimab S309 (19.1 Å) (Figure 2, Table 5), and (3) the antibody LLNL-112 was specifically designed to reduce impact of Omicron RBD’s mutations at positions 446 and 493 on binding to the COV2-2130 (14, 19), (4) the analysis of spike protein sequences reported in GISAID database indicates that mutation Q493R is highly dependent on the presence of H655Y (indeed, in GISAID as of 26/03/2023 there are 6,480,423 sequences with H655Y, 3,745,337 sequences with Q493R, from which 3,742,474 sequences with both mutations which comprise 99.92% cases). In summary, protein structural modeling results support the hypothesis that the detected mutations are in fact antibody escape mutations.

The lack of mAb neutralization displayed by the UP combined with the lack of genetic diversity detected starkly contrasts with the neutralization capability and escape mutants generated by virus stock generated using antiserum selection pressure. One advantage of using viral stock with a very limited passage history, as represented by the UP stock, is the ability to compare assay results between studies. Currently, low passaged stock virus is available from readily obtainable sources such as BEI Resources repository, and this allows many different labs to use the same viral stock. This study indicates that relatively few passages under antisera selection are required to produce mutations that are consistently associated with antibody escape. The antiserum used in this study is comprised of pooled human serum samples from Pfizer vaccinees and is available as reference reagent from BEI Resources, thus providing resources for laboratories to test antiserum selection using reagents from the same source (13). This antiserum reagent was developed specifically as a general reference reagent to be used by different laboratories across the globe for comparison of assay results, and data from this study shows the value of this resource.

## 5. Conclusions

Deep sequencing of COVID-positive clinical samples obtained from immunocompromised patients show the impact of a sub-neutralizing antibody response on the diversity of SARS-CoV-2 intrahost viral populations (1, 44–46) and the emergence of variant genotypes. Even in immunocompetent hosts that mount a rapid and robust immune response to infection to RNA viruses such SARS-CoV-2, the large viral population size combined with the high genome replication error rate, both which characterize RNA viruses, enable evolution of a diverse mutant spectrum (47). As most recently illustrated with SARS-CoV-2, this rampant generation of new genotypes within a host may enable these viruses to adapt to host immunity and occasionally jump to host new species.

The passage protocol used in this study explored the relative impact of passage in multiple cell lines followed by passage under antiserum selection pressure on the generation of variant genotypes and the emergence of mAb escape mutations. Although relatively few new mutations emerged after four passages in three different cell lines, passage with or without antiserum selection pressure rapidly induced distinct sets of mutations. However, only viral inoculum derived from antiserum selection was able to escape neutralization of three different mAbs. Importantly, the viral stock that was not passaged under selection pressure (cell line or antiserum) was not able to produce escape mutants. These data indicate that using viral stock that has been generated under types of selection pressure that reflect the natural environment of the virus may yield more biologically relevant outcomes of phenotypic assays.

## Acknowledgments

This work was performed under the auspices of the U.S. Department of Energy by Lawrence Livermore National Laboratory under Contract DE-AC52-07NA27344. The following reagent was obtained through BEI Resources, NIAID, NIH: Pooled Human Serum Sample, Pfizer Vaccine, NRH-17727. VSV-SARS-CoV-2-S virus and Vero-TMPRSS2 cells received from Sean Whelan’s laboratory (Univ Washington). Antibody LLNL-112 was prepared by Bonnee Rubinfield.

## Supporting Information

Figure S1: Sequence read coverage depth for sequences derived from neutralization assay.; Figure S2: Neutralization assay, GFP expression results.; Figure S3: Comparison of epitope regions for three SARS-CoV-2 neutralizing antibodies and the hACE2 receptor binding motif (RBM).; Table S1: Mutation distribution among samples passaged with or without antibody selection pressure.

