## Supplementary material for "Differential laboratory passaging of SARS-CoV-2 viral stocks impacts the in vitro assessment of neutralizing antibodies": Supplumental Data

Supplemental Figure and Table Data

A

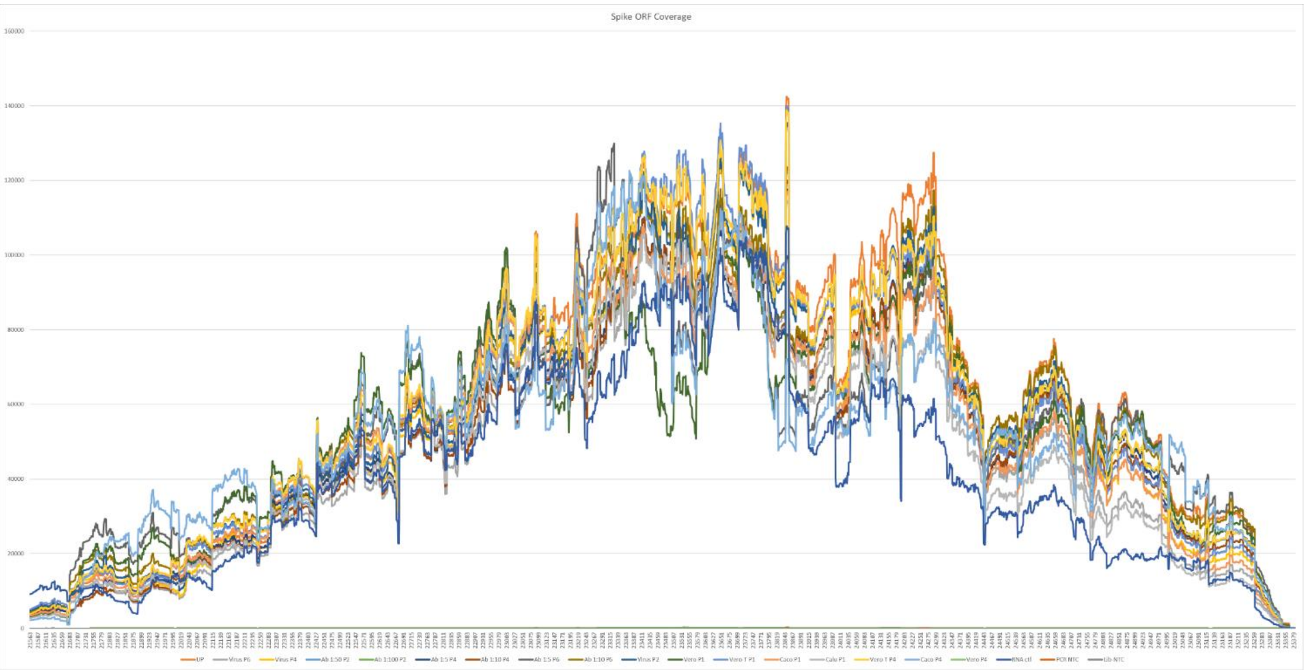

B

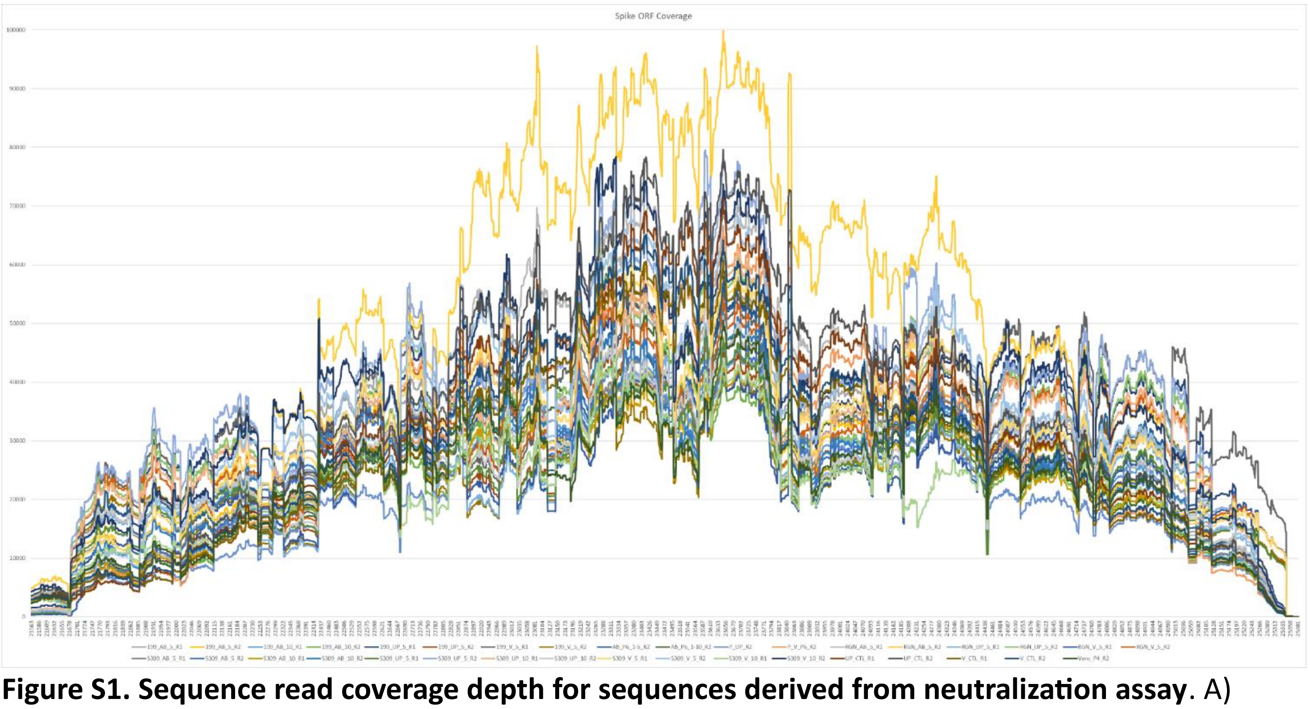

**Figure S1. Sequence read coverage depth for sequences derived from neutralization assay**. A) Sequencing Run 1. Note, NTC yielded just 11 mapped reads. B) Sequencing Run 2. No reads mapped for the NTC.

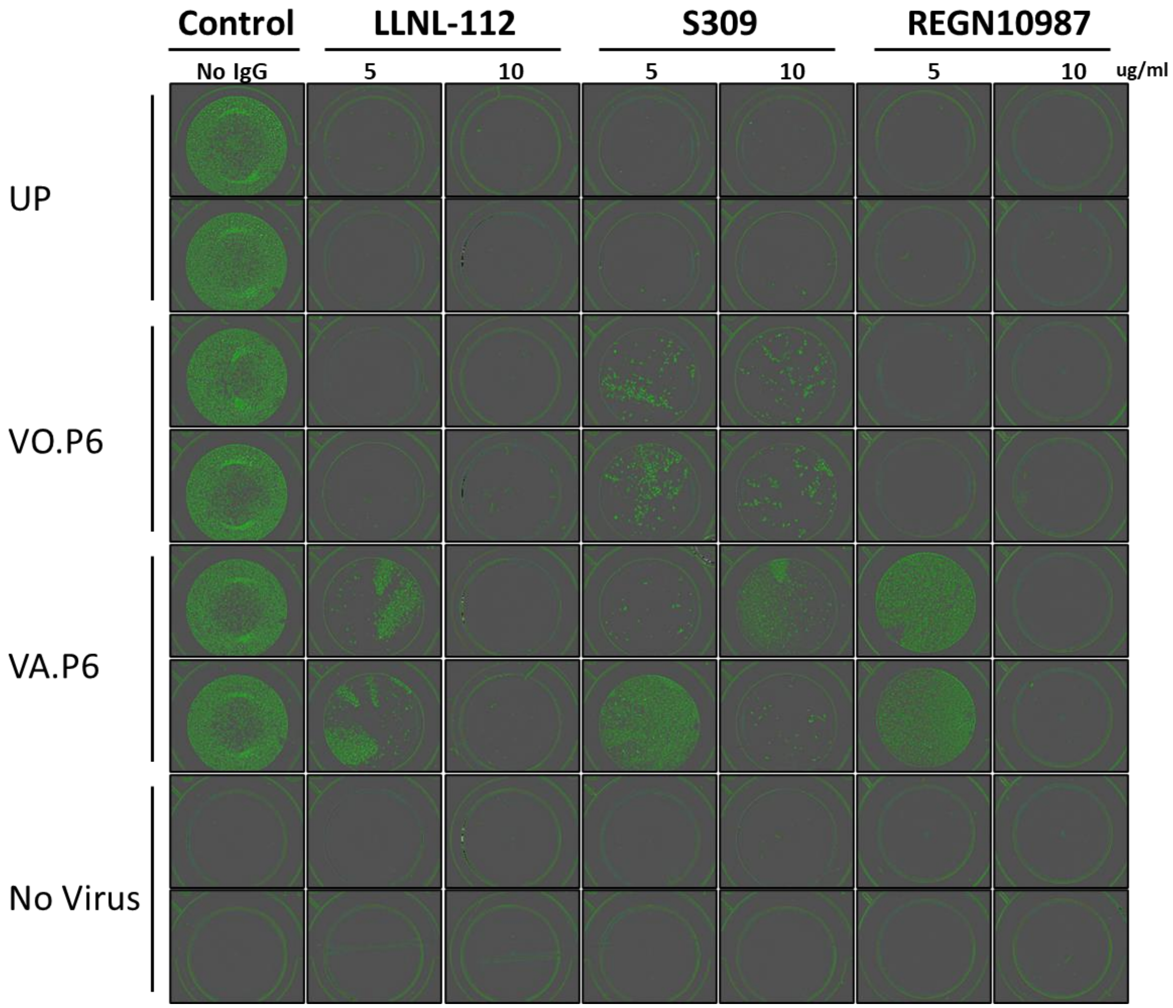

**Figure S2. Neutralization assay, GFP expression results**.

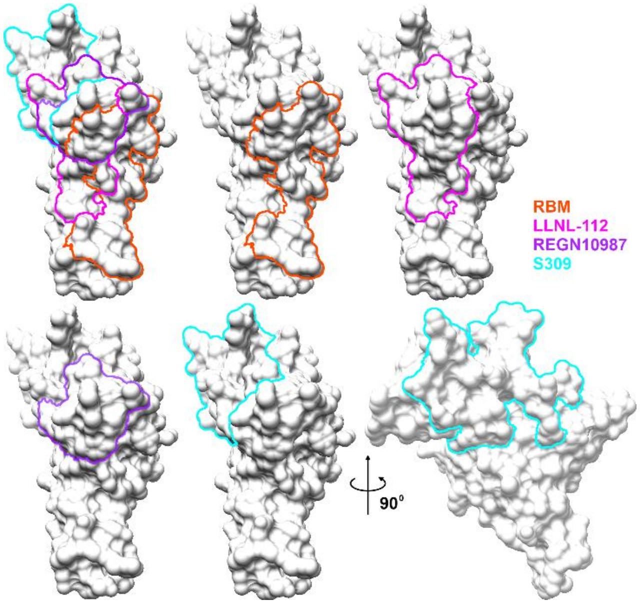

**Figure S3. Comparison of epitope regions for three SARS-CoV-2 neutralizing antibodies and the hACE2 receptor binding motif (RBM).** The epitopes are in close proximity (S309) or overlapping (REGN10987 and LLNL-122) the RBM, which likely accounts for the neutralization effects of these antibodies. The epitope regions and the RBM are outlined on the molecular surface of the receptor binding domain (RBD) of SARS-CoV-2 SPIKE protein. The outlined regions enclose all atoms in any residue that has any atom within 5 Å of any atom of the corresponding antibody and contributes to the molecular surface. The RBD is shown in the same orientation for all panels except for the last panel, which is oriented to show a more complete view of the s309 epitope. This view is 90° around the vertical axis with the right side of the molecule rotated away and the left rotated toward the viewer. This figure was generated using Chimera and GIMP.

**Table S1. Mutation distribution among samples passaged with or without antibody selection pressure.** The frequency of detection (%) of the mutations detected at >1% in antisera selection samples and in samples from the neutralization assay are shown for comparison. Data from virus only passaged samples (no antibody selection) are shown in blue cells and data from virus and antiserum passaged samples are in yellow shaded cells. Top row cells shaded grey signify columns with data from both virus only and antibody selection.

| | Antiserum or mAb (ng) | V47I | L48S | H66R | G72R | T76I | T167I | L242P | H245Y | S247N | Y248N | E340A | R346S | K378E | L390F | K444N | K444T | G446V | T478K | Q493R | I569S | H655Y | E661G | L945F | L977F | N978K | S982L | D985G | V991A | S1003I | I1081V | R1091H | I1216V |
|---|---|---|---|---|---|---|---|---|---|---|---|---|---|---|---|---|---|---|---|---|---|---|---|---|---|---|---|---|---|---|---|---|---|
| **CELL PASSAGE** | | | | | | | | | | | | | | | | | | | | | | | | | | | | | | | | | |
| **UP** | **None** | | | | | | | | | | | | | | | | | | | | | | | | | | | | | | | | |
| **VERO P4** | **None** | | | | | | | | | | | | | | | | | | | | | | | | | | | | | | | | |
| **VERO-T P4** | **None** | | 1.3 | | | 11.7 | | | | | | | | | | | | | | | | 2.5 | | | | | | | | | 2.0 | | |
| **CACO P4** | **None** | | | | | | | | | | | | | | | | | | | | | | | | | | | | | | | | |
| **ANTISERUM PRESSURE** | | | | | | | | | | | | | | | | | | | | | | | | | | | | | | | | | |
| **P_VO_P6** | **None** | | 1.8 | | | 12.0 | | | | | | | | | | | | | | | 9.7 | 82.0 | | | | | | | 1.3 | | 84.4 | | |
| **VA_P6_1-10** | **Antiserum** | | | 1.4 | | 37.4 | | 1.2 | | 1.6 | 1.6 | | 1.4 | | | | | | | 1.7 | | 74.3 | | | | | | | | | 75.5 | | 2.7 |
| **VA_P6_1-5** | **Anitserum** | | | 10.1 | | 24.0 | | | | | | | | | | | | | 1.5 | | | 47.1 | | | | | | | | | 49.3 | | |
| **NEUT ASSAY** | | | | | | | | | | | | | | | | | | | | | | | | | | | | | | | | | |
| ***mAB LLNL-112*** | | | | | | | | | | | | | | | | | | | | | | | | | | | | | | | | | |
| **LLNL-112_UP_5** | **mAB 199 (5)** | | | | | | | | | | | | | | | | | | | | | | | | | | | | | | | | |
| **LLNL-112_V_5** | **mAB 199 (5)** | | 1.8 | | | 12.9 | | | | | | | | | | | | | | | 8.8 | 83.1 | | | | | | | 1.2 | | 83.2 | | |
| **LLNL-112_AB_5** | **mAB 199 (5)** | | | | | 8.6 | | | | | | | 56.7 | | | | | | | | | 88.9 | | | | | | | | | 89.1 | | |
| **LLNL-112_AB_10** | **mAB 199 (10)** | | | | | 60.9 | | | | | | | | | | 52.2 | | | | 58.4 | | 33.8 | | | | | | | | | 36.0 | | |
| ***mAb REGN10987*** | | | | | | | | | | | | | | | | | | | | | | | | | | | | | | | | | |
| **REGN10987_UP_5** | **REGN10987 (5)** | | | | | | | | | | | | | | | | | | | | | | | | | | | | | | | | |
| **REGN10987_V_5** | **REGN10987 (5)** | | 4.0 | | | 11.6 | | | | | | | | | | | | | | | 9.7 | 83.1 | | | | | 1.1 | | 2.0 | | 83.1 | | |
| **REGN10987_AB_5** | **REGN10987 (5)** | | | | | 20.0 | | | 15.7 | | 17.2 | | | | | 31.2 | 16.0 | 40.8 | | 16.0 | | 80.3 | | | | | | | | | 82.1 | | |
| ***mAb S309*** | | | | | | | | | | | | | | | | | | | | | | | | | | | | | | | | | |
| **S309_UP_5** | **S309 (5)** | | | | | | | | | | | | | | | | | | | | | | 5.8 | 2.5 | | | | | | | | | |
| **S309_UP_10** | **S309 (10)** | | | | 3.3 | | | | | | | | | | | | | | | | | | | | | | | | | | | | |
| **S309_V_5** | **S309 (5)** | | | | | 3.7 | 1.3 | | | | | | | 1.8 | 2.6 | | | | | | 46.6 | 90.0 | | | 18.0 | 5.7 | | 1.1 | | | 90.0 | 2.9 | |
| **S309_V_10** | **S309 (10)** | 1.3 | 5.1 | | 1.1 | 8.3 | | | | | | | | | | | | | | | 41.5 | 89.3 | | | | | 14.0 | | 16.0 | 1.4 | 90.0 | | |
| **S309_AB_5** | **S309 (5)** | | | | | 10.1 | | | | | | 57.1 | 1.1 | | | | | | | | | 87.6 | | | | | | | | | 88.7 | | |
| **S309_AB_10** | **S309 (10)** | | | 1.1 | | 13.6 | | | | 1.5 | 1.2 | 30.0 | | | | | | | | 1.1 | | 83.1 | | | | | | | | | 84.0 | | 2.0 |
